# Mobile Laser Scanner understory characterization: an exploratory study on hazel grouse in Italian Alps

**DOI:** 10.1101/2022.04.26.489487

**Authors:** Marta Galluzzi, Nicola Puletti, Marco Armanini, Roberta Chirichella, Andrea Mustoni

## Abstract

Forest vegetation structure assessment is a time expensive effort with traditional methods. The Mobile Laser Scanner (MLS) technology can greatly speed up field works, achieve detailed quantification of three-dimensional forest structure at detailed resolution and drive forest management to increase the conservation status of forests-specialist bird species. In this study, using Mobile hand-held Laser Scanner (MLS), we calculated a fine-scale vegetation density index (namely the Plant Density Index, PDI) to characterize the vertical structure of forest subcanopy (0-10 m). The collected MLS point clouds were used to estimate the abundance of Potential Hiding Refuges (PHR) for the hazel grouse (*Tetrastes bonasia*), a sedentary bird extremely sensitive to forest structure and composition. The study was carried out in 10 plots located in the Adamello Brenta Geopark (Southern Alps, Italy). The species was detected in 8 out of 18 transects in an uneven-aged spruce forest with a discontinuous tree cover. The PDI decreases as the height increases, showing greater value in the shrub and herbaceous layer while the upper values are represented by trees stems, and branches. Visibility analysis of lower understory, highlighted PHR mean value of 73.2% (sd = 9.2%). In our area, PDI and PHR revealed that the environmental factors for hazel grouse occurrence are forests with open habitats, understory vegetation, and good hiding opportunities. Our study is the first application that uses MLS derived parameters to describe the ecological niche of a grouse and we presented the surveyed area as “case report” of hazel grouse habitat.

## 1 Introduction

The hazel grouse (*Tetrastes bonasia*) is a forests-specialist bird that spreads in both lowlands and mountainous regions across Eurasia (Matysek et al., 2020). It is sensitive to forest structure and composition, which are a crucial factor for its distribution and abundance since its sedentary, forest-specialist behavior (Kortmann et al., 2018; Matysek et al., 2020). Hazel grouse prefers large coniferous and mixed forest but can also inhabit early succession stages, as well as small regeneration areas embedded in old-growth forests (Zellweger et al., 2014; Matysek et al., 2020). Its habitat is represented by highly heterogeneous forest conditions due to local disturbance (e.g. wind-throws, snow movements) and management practices oriented to favor multi-layer forest strata in terms of tree and shrubs (Schaublin and Bollmann, 2011; Zellweger et al., 2014).

Vegetation structure influences dynamics of habitat use and hazel grouse forages across life-stages, and different plants including trees, shrubs, and herbs are required to survive different phenological seasons (Ludwig and Klaus, 2017; Zellweger et al., 2014; Matysek et al., 2018). Hence, forest management that leads to habitat structure simplification and forest fragmentation is one of the main threat factors for this species (Mathys et al., 2006; Łukasz Kajtoch et al., 2012; Seibold et al., 2013; European Commission, 2021).

Hazel grouse European population has decreased in the last decades (Storch, 2000; BirdLife International, 2021) particularly in central and Eastern European countries (Rhim, 2006; Matysek et al., 2020). Therefore, it is listed in Annex I of the European Birds Directive (European Commission, 2021). The geographical range of hazel grouse in Italy is limited to the East Alps (BirdLife International, 2021) where trend and size population is difficult to determine, due to the lack of constant monitoring.

Forest-dwelling grouse are highly dependent on forest and vegetation structure (Melin et al., 2016) and several studies have pointed out this relationship with both field and remotely sensed data. Lidar remote sensing has been widely used in wildlife habitat mapping and species distribution modeling due to its capacity to accurately measure three-dimensional vegetation structure (Zellweger et al., 2014; Guo et al., 2017; Kortmann et al., 2018) and both Aerial Laser Scanning (ALS) and static Terrestrial Laser Scanning (TLS) are effective at quantifying components of forest structural diversity and biomass (LaRue et al., 2020; Li et al., 2021), while the use of Mobile Laser Scanner (MLS) is still rare or lacking.

There is a large body of literature concerning the relationship between ALS structural metrics and habitat characterization (Hinsley et al., 2002; BRADBURY et al., 2005; Graf et al., 2009; Melin et al., 2016; Zellweger et al., 2014; Vries et al., 2021).

Using ALS surveys, for example,Melin et al. (2016) showed that species favors forest with dense shrub and heterogeneous canopy, while Zellweger et al. (2014) found that horizontal and vertical canopy height heterogeneity, ground vegetation cover, and basal branches trees are the driver factors of hazel grouse habitat. Besides,Bae et al. (2014) revealed that vertically well-structured forest stands and horizontally mixed successional vegetation stages were key factors of hazel grouse habitat. In addition Mathys et al. (2006) highlighted the influences of forest edge density, the cover of shrub and herb layer on species distribution through satellite images. Recent studies have found a positive relationship between semi-open forest stands with dense understory with hazel grouse presence, while Kortmann et al. (2018) provide a complete review of forest structure attribute associated with hazel grouse in European forest. However, it has been shown that data derived from ALS can influence the ability to access sub-canopy elements due to instrumentation specifications, such as point density and canopy penetration (LaRue et al., 2020).

Static TLS has been only recently used to characterize habitat and ecological niches. For example, TLS was used to investigate the relationship between bat activity and forest structure (Muller et al., 2013; Blakey et al., 2017), to discriminate the nests of black-nest swiftlets from roosting bats in high, inaccessible locations in a cave (McFarlane et al., 2015), while other studies focus on the effects of woodland structure on deer abundance and distribution in the UK (Eichhorn et al., 2017). On the contrary, MLS ecology applications are still lacking (Vierling et al., 2008) and and there are few applications (Nicola et al., 2021).

Considering its higher flexibility for forest field measurements, in this study we capture detailed information on spatial attributes of vegetation presence in forest subcanopy (0-10 m), using MLS data under a voxel-based approach, on hazel grouse forest habitat inside the Adamello Brenta Geopark, Italian Alps. In particular, we aimed to (i) propose a MLS-based approach to define subcanopy characteristics of vegetation of hazel grouse habitat, (ii) estimate the fine-scale vegetation density using the Plant Density Index (PDI) presented in this paper to characterize the vertical structure of forest subcanopy.

Recent ecological studies based on viewshed ecology have used TLS data to understand and quantify the effect of vegetation on animal visibility (Lecigne et al., 2020). Based on these studies TLS allows ecologists to investigate vegetation structure and quantify patterns across heterogeneous and complex habitats and to move beyond two-dimensional measures. The last objective was then to quantify Potential Hiding Refugees (PHR) provided by vegetation structure in the first meters from the ground, an important goal to increase the conservation status of forest grouse.

## 2 Materials and methods

### 2.1 Birds data collection

The study was conducted in the Adamello Brenta Geopark (46°7’17”N, 10°45’8”E) in the south-western part of the Trentino-Alto Adige Region, Southern Alps, Italy (Figure 1). Thanks to the protection led by the Nature Park, the presence of the hazel grouse has been always confirmed since the establishment of the Park in 1988.

**Figure 1:**
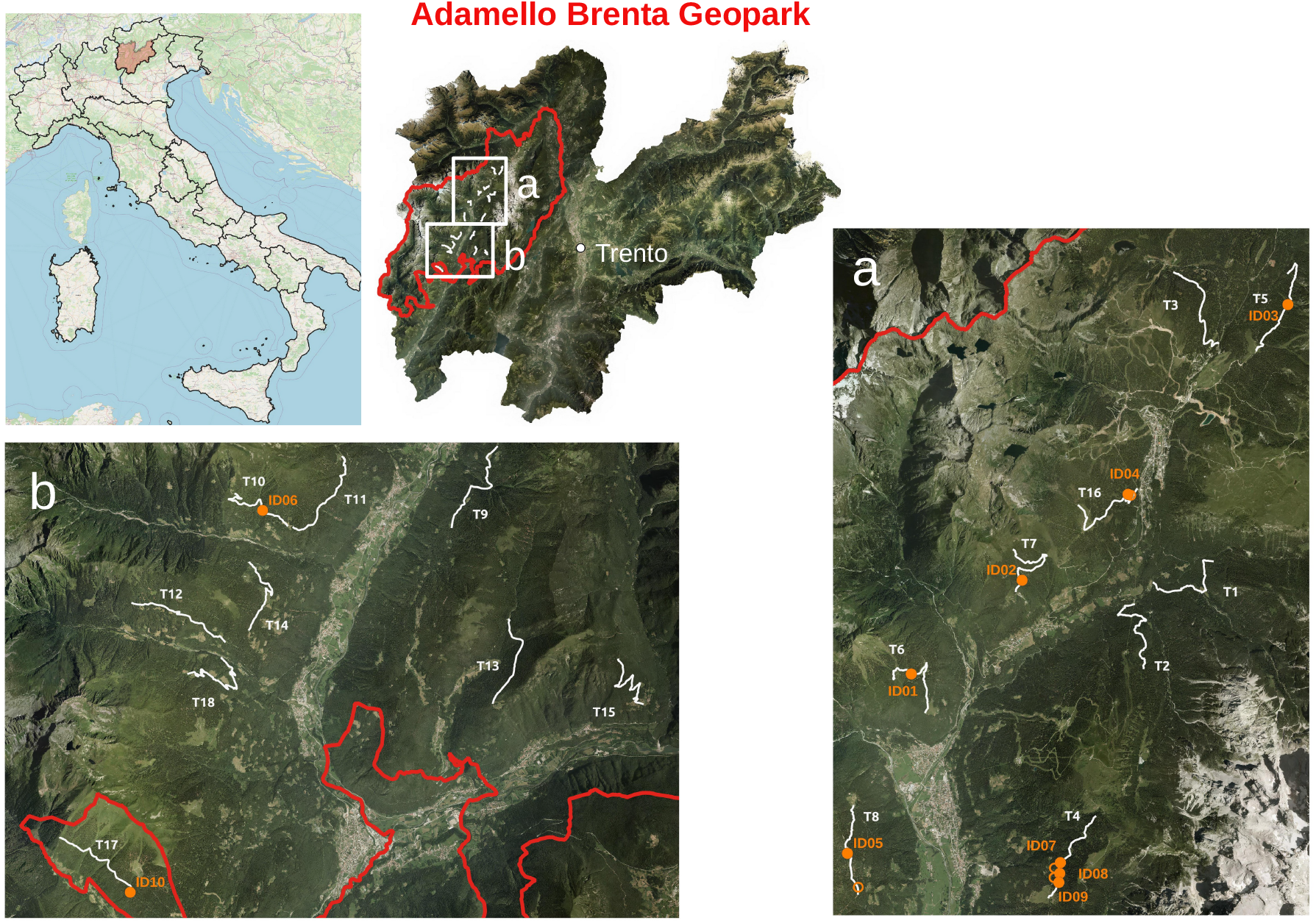
Study area, transects locations and presence track of hazel grouse in the Adamello Brenta Geopark (46°7’17”N, 10°45’8”E; south-western part of the Trentino-Alto Adige Region, Southern Alps, Italy). White lines: transect visited in spring 2021. Orange dot: MLS survey position; Orange circle: hazel grouse tracks where it was not possible to make the surveys with MLS.

Hazel grouse presence was surveyed across potential occurrence places based on previous studies (Mustoni et al., 2008) were 18 transects (ranging from 1100 to 1800 m a.s.l.) were visited twice in spring 2021 during a peak call period (April and May) by using MP3 speakers to play imitation of species. Because of low responses during midday, the census was mainly performed during the morning and evenings and only in good weather conditions (Swenson, 1991; Matysek et al., 2020). Basing on the assumption that playback favors response call on a circle of 100 m radius (Gagliardi and Tosi, 2012), hazel grouse presence was checked every 200-250 m to avoid double counts. In each playback point, the observer did two minutes of listening, and the playback was repeated twice interspersed with two minutes of listening (Swenson, 1991). The species presence was verified by additional checking and searching for tracks such as excrement. A total of 45.78 Km along the transect and 254 playback points were surveyed twice.

### 2.2 MLS data

The hazel grouse forest habitat, from the ground up to 10 m, were described by two different forest stand traits defined using MLS data: (1) the Plant Density Index (PDI) to estimate vegetation density at different point heights (Puletti et al., 2021), and (2) the percentage of Potential Hiding Refugees (PHR) assessed using `viewshed3D` R package (Lecigne et al., 2020).

#### 2.2.1 Data collection and preprocessing

MLS data were acquired between 14 and 17 of June 2021 over the 10 sampling points where hazel grouse was recorded. Round each sampling point, a circle of about 15-20 meters radius was measured using a GeoSLAM ZEB-REVO (GeoSLAM Ltd, Ruddington, England) lightweight mobile laser scanner. The 10 obtained point clouds have been firstly converted to LAS files using the GeoSLAM Hub (GeoSLAM Ltd, Ruddington, England) proprietary software and then clipped on a square of 20 m inside (see (Puletti et al., 2021) for details). The acquired MLS point cloud was normalized using the tlsNormalize function of the `TreeLS` R package (de Conto et al., 2017). The entire dataset is available at https://zenodo.org/record/5653007#.YYjsYp7MKnU (Puletti and Galluzzi, 2021)

#### 2.2.2 Voxelization

To characterize vegetation from MLS data, the production of a 3D matrix of voxels from an MLS point cloud (i.e. voxelization) is commonly used to simplify the subsequent analysis (Puletti et al., 2021). The resulting 3D map can be further analyzed to assess different characteristics, such as, for example, gap fraction, plant density profiles, or Leaf Area Index (Grau et al., 2017) over the whole forest stand. In this study, we used a voxel-grid approach applied to arrange the 3D-space (a box of 20 m x 20 m x 10 m) in voxels of 0.5 m x 0.5 m x 0.25 m in x, y, z dimensions, respectively.

This step was performed using the `crossing3dforest` (Puletti and Galluzzi, 2021) R package.

#### 2.2.3 Plant Density Index

Starting from a voxel-grid approach, in order to define structural three-dimensional characteristics of vegetation in the forest subcanopy layer, a Plant Density Index (PDI) (Eq.1) was computed:

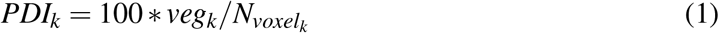

where *veg_k_* is the number of vegetated voxels in the *k_th_* voxel height and *N_voxel_k__* is the total number of voxels in the same voxel height.

Firstly, *PDI_k_* describes vegetation density profiles for each *k_th_* voxel height (0.25 m). *PDI_k_* values are then aggregated considering lower understory (LU, 0-5 m) and upper understory (UU,5-10 m) layers.

#### 2.2.4 viewshed3D application

The lower understory (LU, 0-5 m) voxel-grid were processed by *viewsheds()* function of `viewshed3D` R package to identify potential hiding refugees. This function allows to quantify the relative visibility of the environment by combining viewsheds from multiple, user-selected viewpoints into a single variable that estimates a cumulative viewshed (Lecigne et al., 2020).

For each plot, we computed visibility from 100 viewpoints systematically distributed on the lower understory (LU, 0-5 m) stratum. To represent the potential viewpoints of a fox (i.e. one of the common hazel grouse predators), the viewpoint heights were set at 1 m above the ground. Potential Hiding Refugees (PHR) were then identified as voxels that were visible from an arbitrarily selected threshold of <5% of all sample viewpoints.

### 2.3 Understory characterization

In each sampling plot, the most important species belonging to tree, shrub, herbaceous, and regeneration layer were recorded. For each species, the Braun-Blanquet cover-abundance scale (Braun-Blanquet, 1932) and the main height were noted to better characterize hazel grouse habitat.

## 3 Results

### 3.1 Hazel grouse occurrence

Hazel Grouse was detected in 8 out of 18 transects (Figure 1) for a total of 14 records (6 playback responses and 8 tracks). All hazel grouse signs were found in an uneven-aged spruce forest with a discontinuous tree cover due to forest management and in plot ID03 due to avalanche.

Forest regeneration, composed by *Picea abies, Fagus Sylvatica, Sorbus aucuparia*, was found both aggregated in clusters or randomly distributed, except for one plot where no forest regeneration was found.

The surveys were carried out in 10 points out of 14 signs of presence found since 1 sign (located in T16 Figure 1) was a double count and in 3 plots (located in T8 and T4 Figure 1) the local conditions did not allow access with MLS. The list of surveyed plots and a summary of percentage cover range of tree, shrub and herbaceous layer and regeneration type is presented in Table 1. Secondary tree species such as *Fagus sylvatica* and *Larix decidua* were found. *Fagus sylvatica*, *Sorbus aucuparia* and *Laburnum anagyroides* were the most frequent species in the shrub layer, while *Vaccinum myrtillus, Brachipodium* sp., *Rubus idaeus* and *Dryopetris filix-mas* in the herbaceous layer. See Table 2.

**Table 1:** Summary of surveyed plots and percentage cover range of tree (>3 m), shrub (3-0.5 m) and herbaceous (0-0.5 m) layer.

| Plot ID | Transect | Point type | Tree cover% | Shrubs cover% | Herbac. Cover% | Regeneration type |
| --- | --- | --- | --- | --- | --- | --- |
| ID01 | T6 | excrement | 20-40 | 0-20 | 80-100 | sporadic |
| ID02 | T7 | excrement | 40-60 | 0-20 | 40-60 | sporadic |
| ID03 | T5 | excrement | 20-40 | 0-20 | 20-40 | sporadic |
| ID04 | T16 | call | 20-40 | 0-20 | 40-60 | group |
| ID05 | T8 | call | 20-40 | 40-60 | 40-60 | sporadic |
| ID06 | T10 | excrement | 60-80 | 20-40 | 0-20 | group |
| ID07 | T4 | excrement | 40-60 | 40-60 | 40-60 | group |
| ID08 | T4 | excrement | 20-40 | 40-60 | 40-60 | group |
| ID09 | T4 | excrement | 20-40 | 20-40 | 40-60 | absent |
| ID10 | T17 | call | 60-80 | 20-40 | 0-20 | group |

**Table 2:**
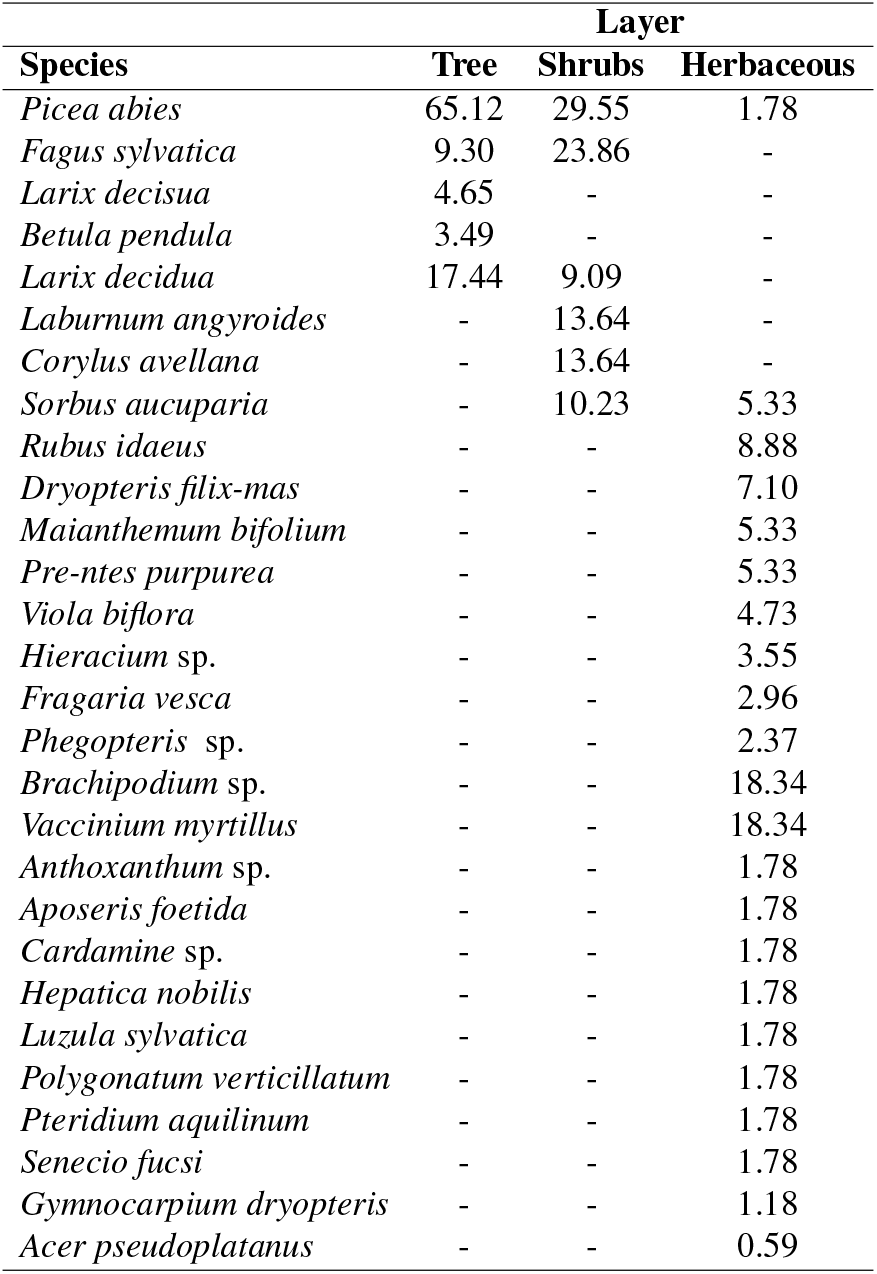
Percentage abundance of species in the tree, shrub, herbaceous layers.

### 3.2 Understory characteristics

The proposed Plant Density Index (PDI, Figure 2) shows the vertical profile of vegetation density from 0 to 10 meters. As expected the PDI decreases with height, showing greater value in the shrub and herbaceous layer, while the upper values are represented by tree stems and lower branches.

**Figure 2:**
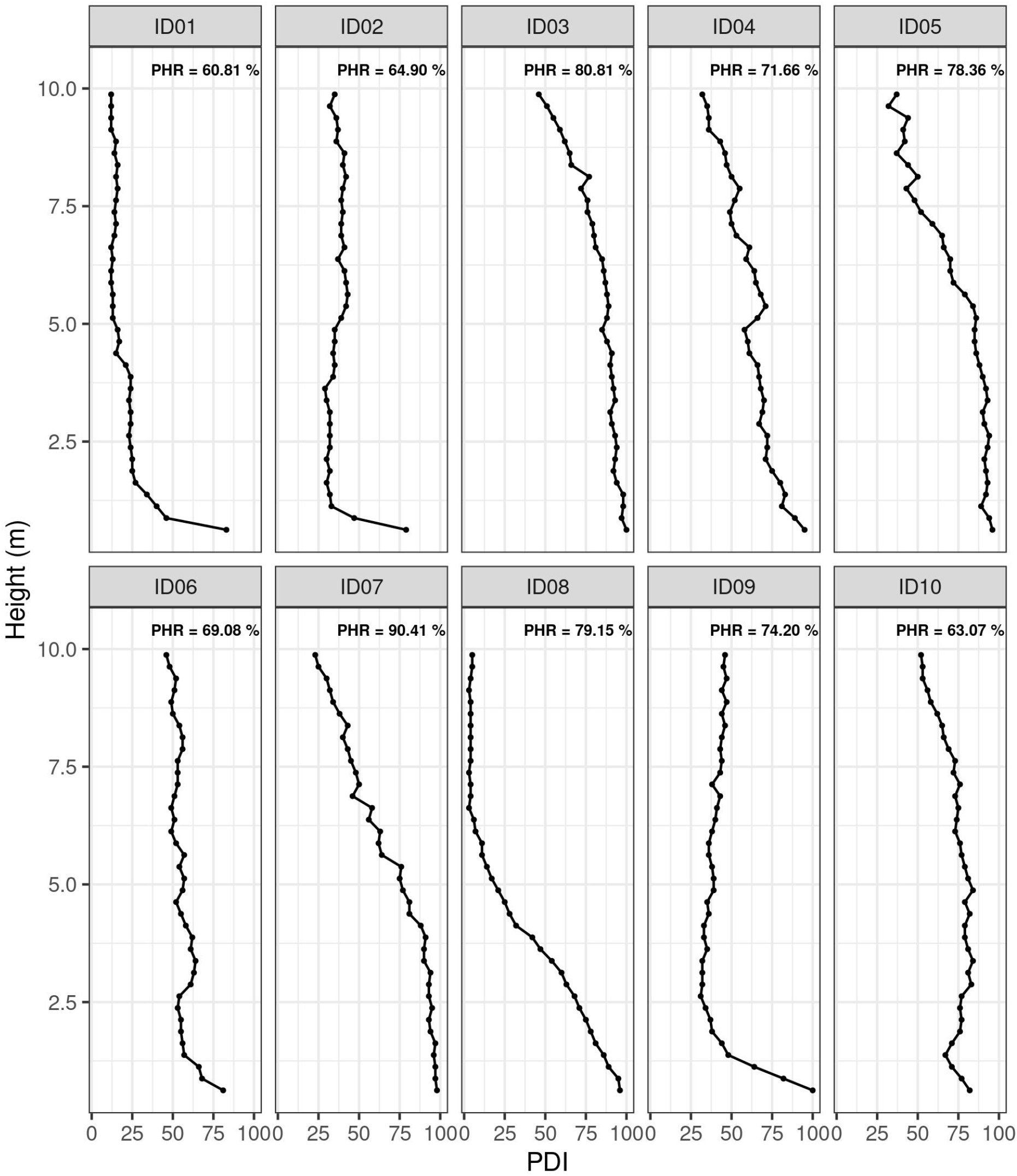
Subcanopy Plant Density Index (PDI) profile by plot (0-10 m) and Potential Hiding Refuges (PHRs) values of lower undestory (LU: 0-5m), derived from visibility analysis computed by `viewshed3D` R package.

Figure 2 shows the comparison of PDI between plots. All the plots showed vegetation density greater than 60% in the first meter from the ground due to the presence of abundant and dense *Brachypodium sp.* and *Vaccinium myrtillus*. Plots ID01, ID02, ID09 have PDI close to 25-30% at 2-3 m, which reveals not very dense vegetation, and thus the presence of gaps, in the shrub layer (Table 1). In the same height range, plot ID06 has density values close to 50% composed by *Fagus Sylvatica* and great stems of *Picea abies*, while plots ID04 and ID10 have PDI value close to 75% represented by dense shrubs (*Fagus Sylvatica, Laburnum angyroides, Corylus avellana*) and tree subject with many branches. Plot ID08, where no shrubs are present has density values close to 75% at 2-3 m, mainly composed by a dense regeneration group of *Picea abies*, with upper vegetation composed by few big trees (Table 1).

Plots ID03, ID05, ID07 have the greatest value of vegetation density (> 80%) in the lower understory, due to the combined effect of denser shrubs, tree subject with many branches, and a dense regeneration group of *Picea abies*. With canopy density reductions, all the plots showed a PDI decrease in the layers included in the range 5-10 m. Plots ID01 and ID08 have PDI <25% above 5 m: in these plots, only few stems (less than five) of *Picea abies* occupy the available space in the tree layer (Table 1). In the other plots, upper understory density values range from 25% to 50%.

Viewshed 3D analysis of lower understory (LU, 0-5 m), highlighted PHR mean value of 73.2%(sd = 9.2%), with the highest value in plot ID07 (PHR = 90.4), while plot ID01 has the lowest value (PHR = 60.8%). PHR values for each plot are represented in Figure 2.

## 4 Discussion

### 4.1 Hazel grouse census

Estimate relative density and presence/absence data are necessary to address many ecological problems (Swenson, 1991). Many birds that are difficult to detect in the field, respond well to playback recordings of their calls, including hazel grouse (Swenson, 1991).Swenson (1991) found hazel grouse respond well both in spring and autumn with a response rate of 80%. A similar response rate was found by Łukasz Kajtoch et al. (2012) in a long-term study (from 2000 to 2010) however the contactability of the species over the years has varied a lot between forest patches (from 16% to 80%). Hazel grouse’s presence in the Adamello Brenta Geopark is historically documented by previous studies (Mustoni et al., 2008) and by wildlife monitoring projects conducted from 2005 to 2012 by Park rangers. However, during this census, the species showed a very low call response rate (2.4%). This may be due to many factors including annual forest patches change by species (Łukasz Kajtoch et al., 2012), climatic condition, and also a decrease in the abundance of the species. The winter before the census was characterized by a very abundant historical snowfall that may have influenced the species behavior and local abundance (Gavrin, 1969; H.H. Klaus, 1982; Saniga, 2002). Due to the low response rate, it was not possible to ascertain the true areas of the absence of the species. Further census replicas are required to assess more detailed species spatial distribution. For these reasons, the present work takes on the character of a “case report” where the potential of MLS in applied ecology was shown.

### 4.2 Hazel grouse habitat

Habitat quality and forest structure are crucial for the distribution of many species of birds and many studies showed that several factors affect hazel grouse occurrence (Łukasz Kajtoch et al., 2012; Matysek et al., 2018). Basing on this exploratory study hazel grouse inhabited uneven-aged coniferous forests and mixed forests mainly composed of spruce and secondary from beech and larch, as reported in other studies (Łukasz Kajtoch et al., 2012). In addition, in the Adamello Brenta Geopark, sites occupied by hazel grouse were characterized by lower tree cover due to forest management or natural disturbances like avalanche a confirmation of what has been observed by Kortmann et al. (2018) where the probability of hazel grouse presence increased with increasing disturbance, and with successions dynamics. Uneven-aged forest with horizontal and vertical cover variability is characterized by greater light incidence on the ground which causes the development of herbaceous and shrub layer.Melin et al. (2016) found hazel grouse occurrence positively correlated with increasing shrub cover (Melin et al., 2016), Aberg et al. (2003) found denser ground and shrub cover associated with the species and Swenson (1991) found that dense vertical vegetation cover is important for hazel grouse survival. The most frequent species in the shrub and herbaceous layer were *Fagus sylvatica, Sorbus aucuparia, Laburnum anagyroides* and *Vaccinum myrtillus, Brachipodium sp., Rubus idaeus* and *Dryopetris filix-mas* respectively. The occurrence of trees in the understory as mountain ash (*Sorbus aucuparia*) is an important source of food as shown in other studies conducted in Poland (Matysek et al., 2018) and southeast Germany (Bae et al., 2014). Recent studies have revealed that the number of tree species in the shrub and herbaceous layer had a significant impact on the probability of hazel grouse occurrence (Matysek et al., 2020). Furthermore, the presence of bilberry is well documented for hazel grouse in spring, autumn, and post-breeding period habitat (Łukasz Kajtoch et al., 2012; Bae et al., 2014; Ludwig and Klaus, 2017; Matysek et al., 2018). Bae et al. (2014) reported that in southeast Germany, the species composition of the shrub layer appears to be of higher importance for habitat suitability than the cover of the shrub layer. More generally, the shrub and herb layers constitute important habitat characteristics due to determining the availability of nutrition and shelter for the birds (Vauhkonen and Imponen, 2016).

Vegetation structure is an important feature of grouse habitats and Lidar data can characterize this structure more precisely than a traditional survey. In a traditional survey, layers cover is subjectively assessed based on 2D percentage classes (Table 1) while the MLS survey objectively assesses the true layers cover taking into account all the components at a certain height in a 3D space (i.e. herbaceous cover plus stems and branches cover). The plots surveyed in this study reveal different vegetation density height profiles (Figure 2). For example, basing on the traditional survey plots ID01, ID02 and ID09 have very low percentage cover (0-20%, Table 1) and no shrubs were present. While basing on MLS data, plots ID01, ID02, and ID09 have PDI close to 25-30% at 2-3 m which represents space occupied by stems and thus highlights the presence of gap along with the vegetation density. The importance of open areas along the canopy cover has been reported by several studies and leads to an increase in the probability of species presence (Luque and Delcros, 2013; Kortmann et al., 2018; Matysek et al., 2018, 2020). In a study conducted in southern Poland by Łukasz Kajtoch et al. (2012), canopy openings were the most important factor for hazel grouse occurrence. As reported by a recent review on habitat requirements of hazel grouse in European forest (Kortmann et al., 2018), the species prefer heterogeneous forest structures with dense herbaceous cover (close to 50%) and dense shrubs. The heterogeneity of herbaceous and shrub layers has been performed by traditional survey or ALS data which have the limit of low pulse density at certain height (Melin et al., 2016). Until now, no detailed study on sub-canopy structure was conducted by using MLS data and this study wanted to explore the potentiality of new technologies applied to ecological studies.

The visibility analysis has proven to be valid and in line with the values of PDI (Equation 1) in the lower understory (LU: 0-5m) (Figure 2). Specifically, plot ID03, with a PHR of 80.81%, has a very dense vertical profile with PDI greater than 75% in the LU. On the contrary, plot ID01 which has the lowest value of PHR (60,81%), also has the lowest values of PDI in the LU (Figure 2). In all plots, PHR has values greater than 60%, which reveal the importance of hiding refuges and thus habitat heterogeneity in hazel grouse occurrence. As previously highlighted by Seibold et al. (2013) predation rates decline with increasing density of near-ground vegetation. More in general simplified forest structures increase the possibility of moving and searching for prey by predators (Matysek et al., 2018). Although it found results in accordance with what was observed, the viewshed approach highlighted aspects that deserve to be discussed. First of all, as well known in forestry applications, the effect of the steep slope on MLS uncertainty in vegetation monitoring. (Barbarella et al., 2017). In this work, the normalization procedure of the MLS points cloud has not been automated for the various plots and each cloud has been normalized properly, a procedure that required expensive work. Secondly, the random sampling of `viewshed` function does not take into account the limitation of edge effect on visibility. To cope with this issue the Digital Terrain Model (DTM) has been expanded, placing the viewpoints even outside the scan area, and further cropped in the area of interest only.

Dense forest floor are important for hazel grouse because they provide a rich food base and possibilities to hide (Matysek et al., 2018) and protection against predator (Melin et al., 2016). Furthermore, the areas studied were characterized by uneven-aged forest with different successions stages and thus by a discontinuous cover. The importance of such structure was found in several studies.Matysek et al. (2020) found open areas such as clearfelling with dead wood and hollows and herbs layer are important both in spring and winter territories and Łukasz Kajtoch et al. (2012) found the presence of clearings and pioneer trees the most important factors for the species. Habitat heterogeneity has a great importance on prey-predators relationship hence, a simplified forest structure increases the possibility of predation (Seibold et al., 2013). The density of herbaceous and shrub layers influence the predation rates which decline in dense understory (Seibold et al., 2013).

Key factors of hazel grouse distribution are habitat structure and landscape heterogeneity (Luque and Delcros, 2013; Matysek et al., 2018). In conservation and management plans, special attention should be paid to habitats and further studies to deepen our knowledge of subcanopy heterogeneity could be help full for forest management and for the protection of the species.

Forest structure is one of the most important drivers for habitat selection and distribution of many bird species (Cody, 1981). Preservation of various forest structures is a key strategy for the protection of endangered species (Łukasz Kajtoch et al., 2012; Zellweger et al., 2014; Matysek et al., 2020). Habitat fragmentation and transformation are the main threat for forest-dwelling birds, such as hazel grouse, leading to heterogeneity loss (Łukasz Kajtoch et al., 2012).

## 5 Conclusion

The ability to continuously evaluate understory vegetation structure enhances our understanding of wildlife habitat selection, ecology and thus improves forest management. In this study, we showed the potentiality of MLS data on habitat suitability of hazel grouse as a “case report”. It is the first application that uses MLS-derived parameters to describe the ecological niche of a grouse. Unfortunately, due to the low response rate of the species, we could not use MLS data to properly model the habitat requirement of the species. Further monitoring campaigns should be implemented to both increase the presence data of hazel grouse and to perform statistical analyses over an adequate number of sampling areas. However, this study made it possible to highlight, although only in descriptive terms, the ability of MLS to capture fine-scale ecological properties, impossible to measure with visual analysis. Specifically, MLS data allowed to define detailed vegetation density values and to quantify the presence of hiding refugees. We thus encourage continuous research in the ecological study as gained from MLS data. Generally, our results show that MLS data will enable conservation managers, especially of protected areas, to assess fine-scale habitat suitability in areas particularly sensitive and important for the conservation of the species. MLS thus offers great potential for effective ecological monitoring and management and conservation of endangered species.

## Acknowledgement

This paper was funded with the contribution of the Italian Ministry of Agricultural, Food, and Forestry Policies (MiPAAF) subproject “Precision Forestry” (AgriDigit program) (DM 36503.7305.2018 of December 2, 2018). We thank Maurizio Cenci, our collegue of CREA research center, for the assistance with MLS surveys.

